## Supplementary Information for "Interpretable Protein-DNA Interactions Captured by Structure-Sequence Optimization"

#### Summary

**1 Details of Molecular Dynamics Simulation S-2**

**2 Comparison of IDEA predictive performance Using HT-SELEX data S-2**

**3 Analysis of examples where IDEA fails to recognize strong DNA binders S-5**

**4 Supplementary Figures S-6**

#### 4 Supplementary Figures

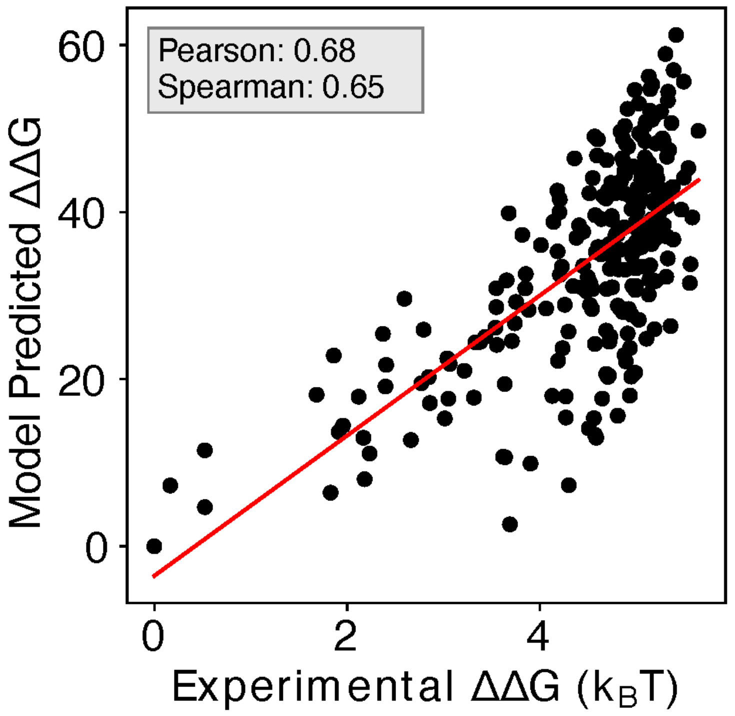


Figure S1: **Including additional human Max-DNA complex structures in IDEA training improves prediction.** Including additional human-associated MAX-DNA complex structures and their associated sequences (PDB IDs: 1HLO, 1NLW, 1NKP) leads to a minor improvement in the correlation between the predicted and experimental binding free energies, with a Pearson correlation coefficient of 0.68 and Spearman's rank correlation coefficient of 0.65. $\Delta\Delta G$ represents the changes in binding free energy relative to the protein binding to the wild-type DNA sequence. The predicted binding free energies are presented in reduced units, as explained in the Methods Section: *Training Protocol*.

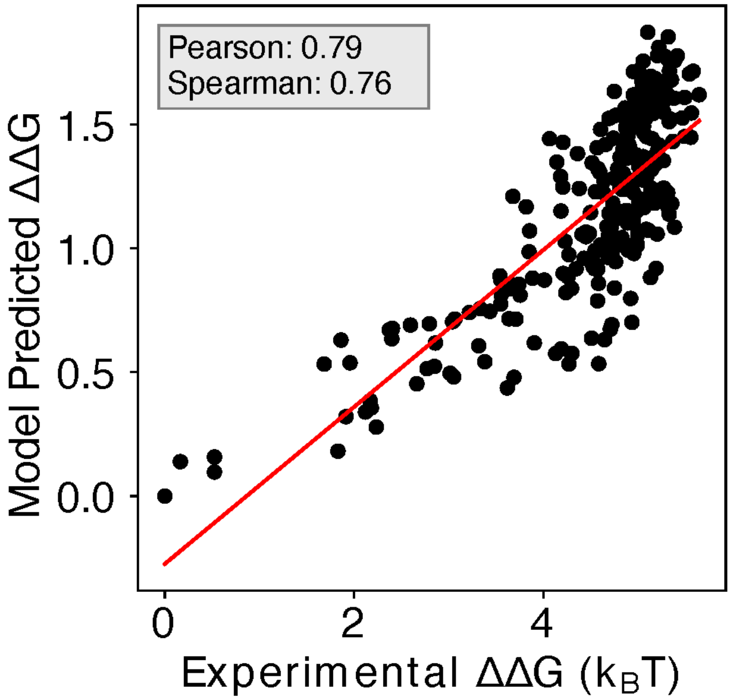


Figure S2: **Enhanced IDEA prediction by integrating SELEX-seq data for MAX transcription factor.** Incorporating SELEX-seq data[S20] into IDEA's training protocol significantly improves its predictive accuracy on MITOMI binding measurements[S21] for the MAX transcription factor, achieving a Pearson correlation coefficient of 0.79 and a Spearman's rank correlation coefficient of 0.76.

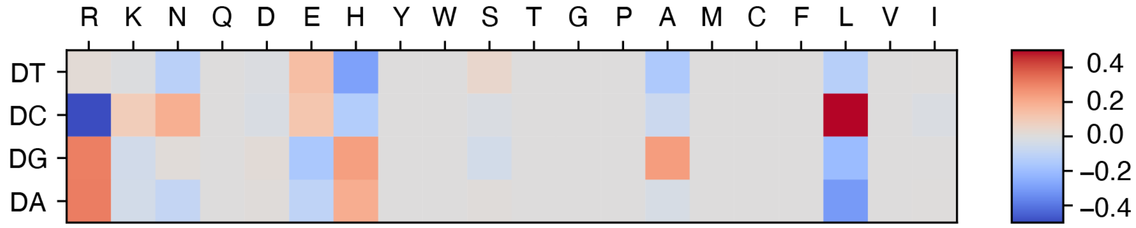


Figure S3: **Refined energy model with SELEX data integration reveals additional physicochemical insights into protein-DNA interactions.** Upon integrating the SELEX data into our model training, we found that the refined energy model reveals additional unfavorable interactions between glutamic acid (E) and deoxycytidine (DC), consistent with their negative charges. The optimized energy model is presented in reduced units, as explained in the Methods Section: *Training Protocol*

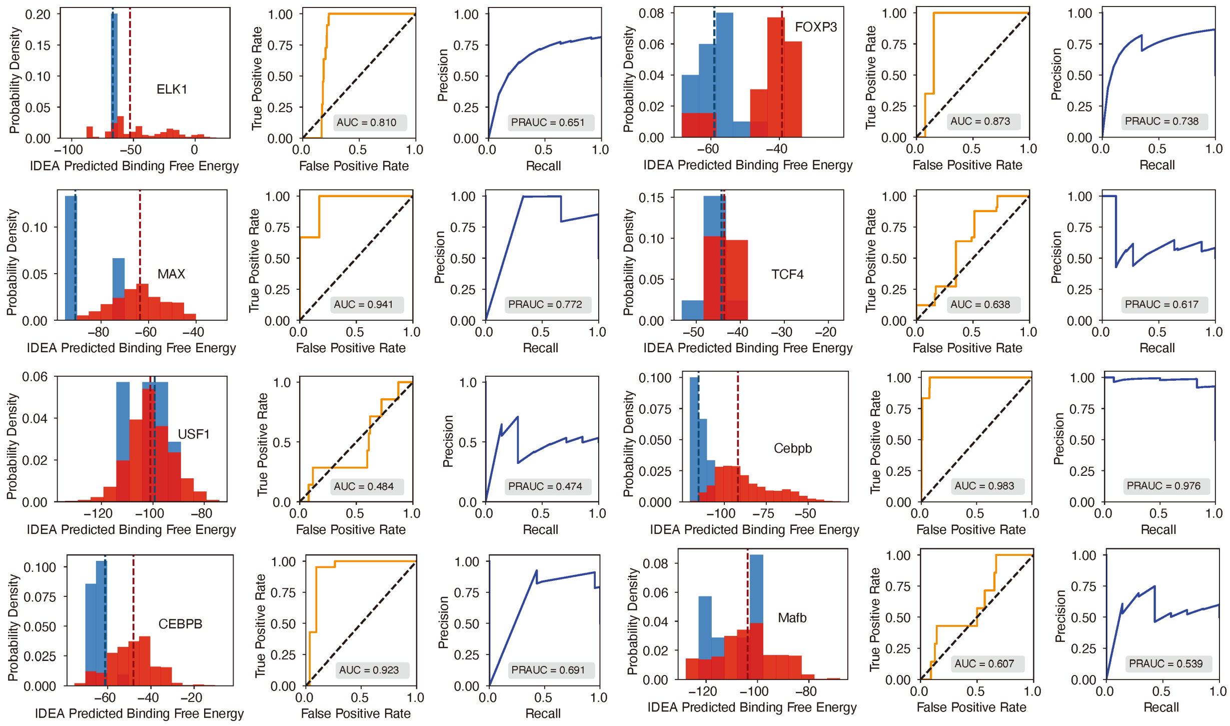


Figure S4: **Evaluation of IDEA's predictive accuracy for distinguishing strong from weak protein-DNA binding interactions (Part 1 of 3).** Probability density distributions of predicted binding free energies for strong (blue) and weak (red) binders identified from the HT-SELEX experiment[S9] are shown across multiple protein families. A clear separation between these distributions indicates the model's effectiveness in identifying high-affinity DNA binders. The darker dashed lines within each distribution correspond to the median binding free energy for strong (blue) and weak (red) binders, respectively. For Mafb, the predicted median binding free energies for strong and weak binders are nearly identical, measured at −103.826 and −103.824, respectively. As a result, the dashed lines almost completely overlap, and only one appears visible in the plot. The Receiver Operating Characteristic (ROC) curve and Precision-Recall (PR) curve adjacent to each density plot provide a quantitative assessment of prediction accuracy, with the Area Under the Curve (AUC) and precision-recall AUC (PRAUC) scores displayed in each panel.

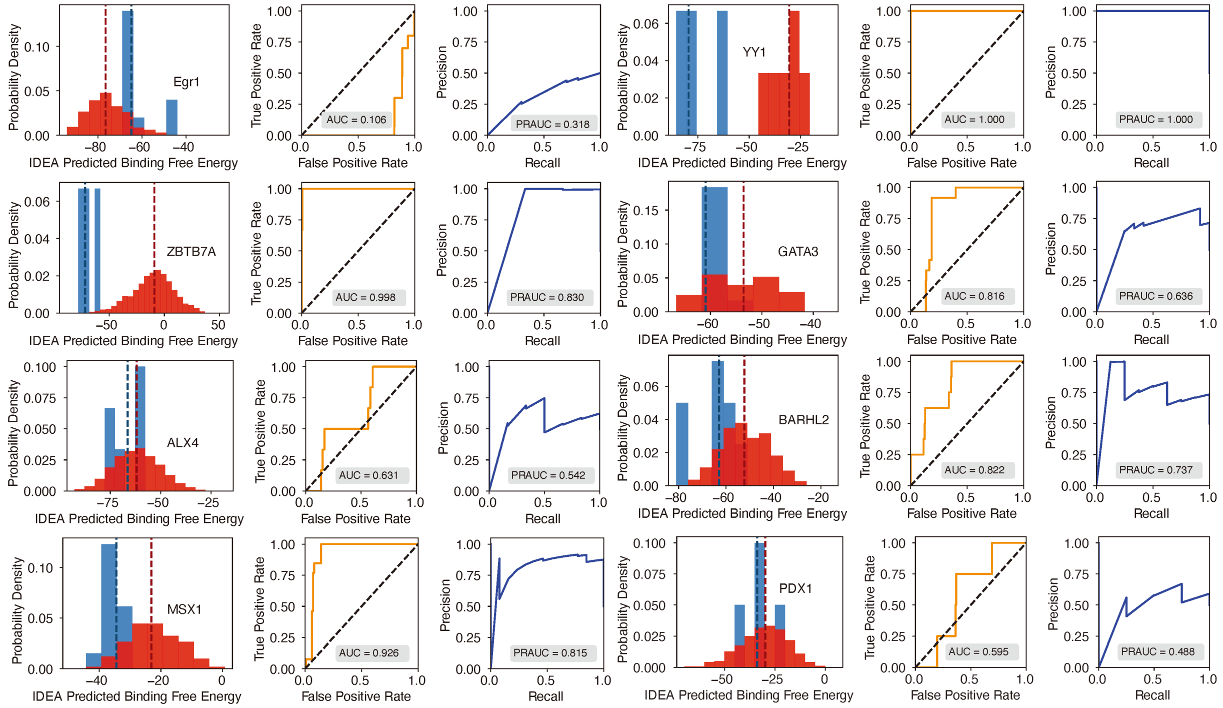


Figure S4: Evaluation of IDEA's predictive accuracy for distinguishing strong from weak protein-DNA binding interactions (Part 2 of 3).

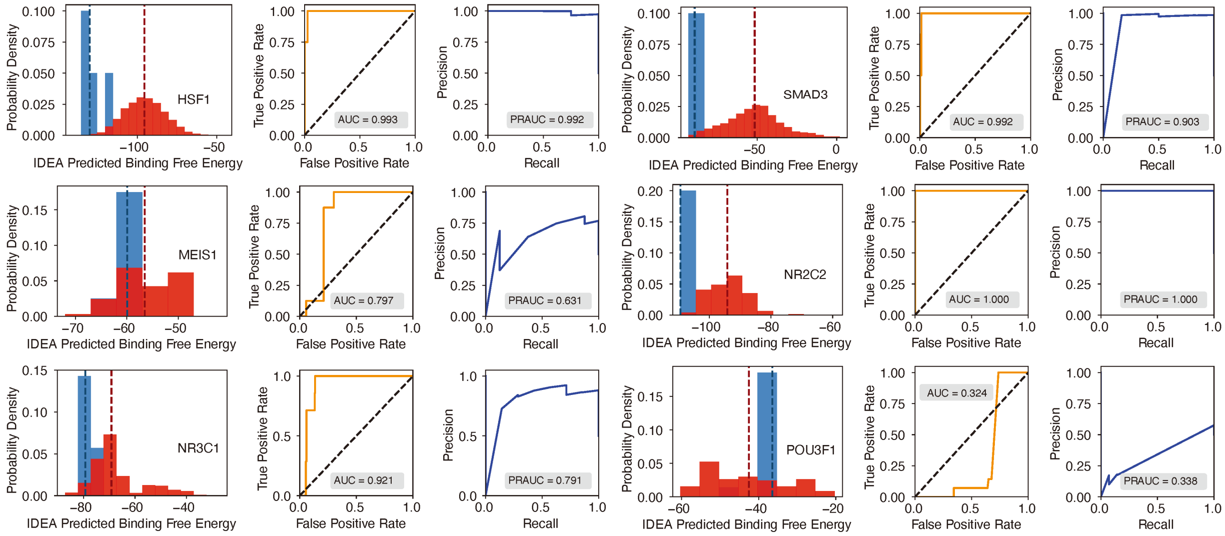


Figure S4: Evaluation of IDEA's predictive accuracy for distinguishing strong from weak protein-DNA binding interactions (Part 3 of 3).

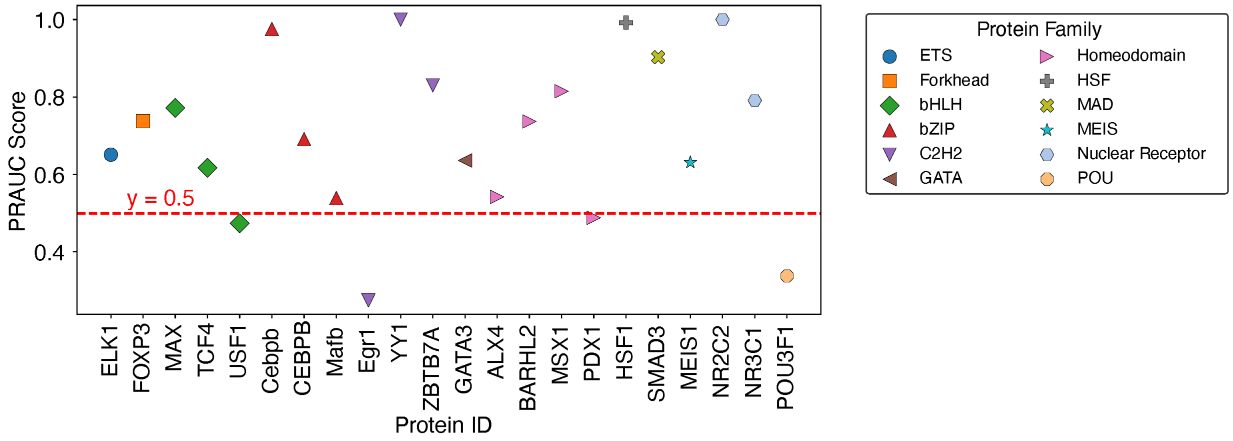


Figure S5: **Summary of balanced PRAUC scores for protein-DNA pairs across 12 protein families.**

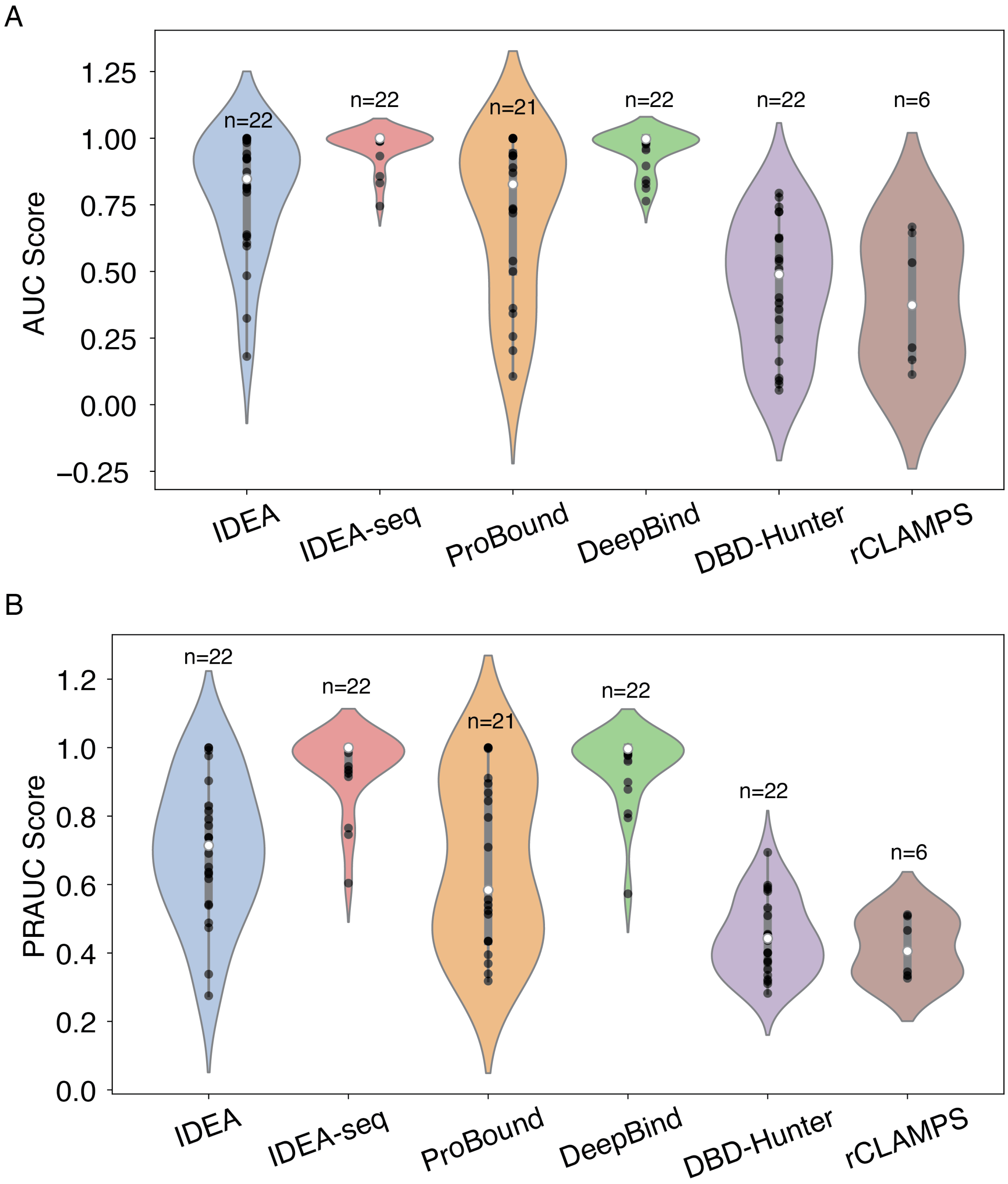


Figure S6: **Performance comparison of IDEA model with other prediction methods.** (A) AUC scores and (B) PRAUC scores for identifying strong binders were evaluated across six different prediction methods: IDEA, IDEA augmented with binding sequences (IDEA-seq), ProBound,[S10] DeepBind,[S11] the general knowledge-based energy model DBD-Hunter,[S17] and family-specific knowledge-based energy model rCLAMPS.[S22] Predictive performances were assessed using HT-SELEX datasets spanning 22 proteins from 12 protein families. Each violin plot displays the distribution of scores for individual targets, with the width indicating score density. The thick grey bar represents the interquartile range (first to third quartiles), and the thin line extends to 1.5 times the interquartile range. Individual data points are depicted as scattered black dots, and the white dot represents the median. Sample sizes ($n$) are labeled above each group.

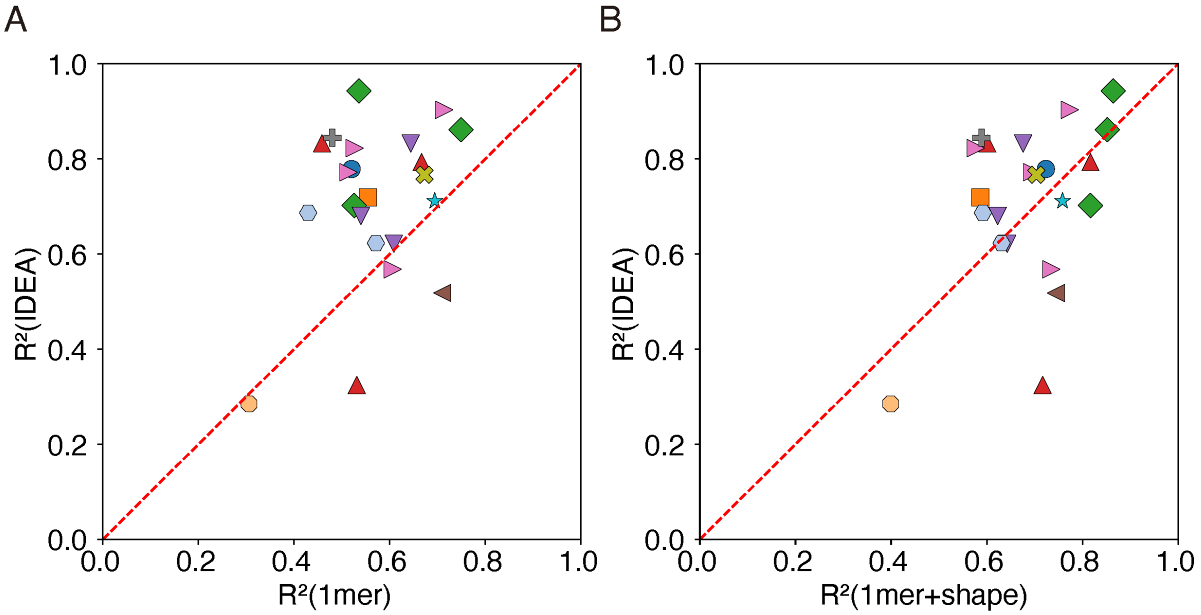


Figure S7: **IDEA outperforms other predictors in cross-validation analysis of protein-DNA binding affinity.** Results from a 10-fold cross-validation indicate that IDEA achieves higher $R^2$ scores for most of the protein-DNA complexes that have available experimentally resolved structures, compared to the 1mer and 1mer+shape methods reported in.[S9] In panel (A), each point represents a protein-DNA complex and compares IDEA's $R^2$ score with that of the 1mer method, while panel (B) compares IDEA's $R^2$ score with the 1mer+shape method. Points are color-coded by protein family (see Figure 2E for family annotations).

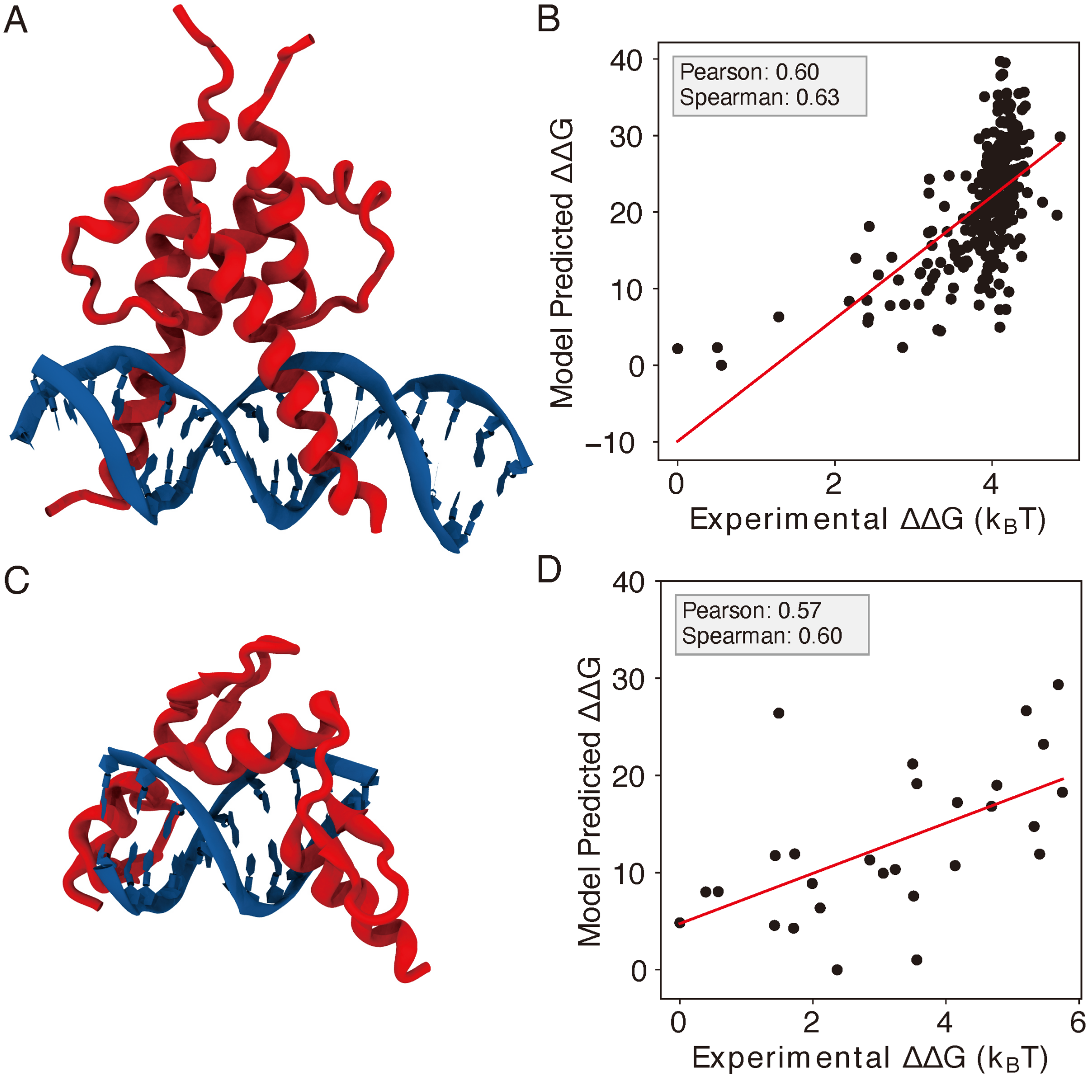


Figure S8: **IDEA correctly predicts the protein-DNA recognition by additional transcription factors.** (A) The 3D structure of basic helix-loop-helix (bHLH) transcription factor PHO4 and its associated DNA (PDB ID: 1A0A). (B) Model-predicted binding free energies of PHO4 correlate with the experimentally determined binding free energies.[S21] (C) The 3D structure of zinc finger protein Zif268 and its associated DNA (PDB ID: 1AAY). (D) Model-predicted binding free energies of Zif268 correlate with the experimentally determined binding free energies.[S18] $\Delta\Delta G$ represents the changes in binding free energy relative to the protein binding to the wild-type DNA sequence.

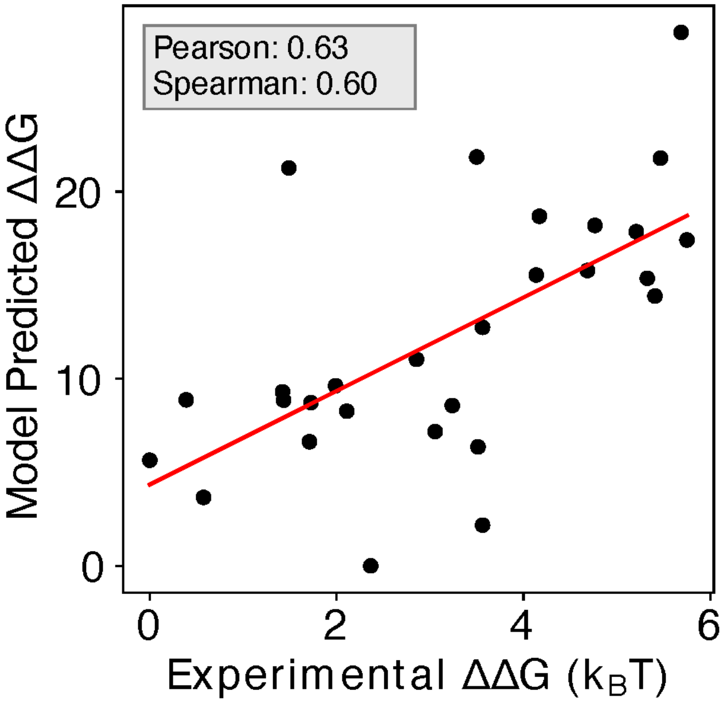


Figure S9: **Enhanced Predictive Accuracy with Inclusion of Zif268 and Related CATH Protein Structures.** Including the structures and associated sequences of 1AAY and other protein-DNA complexes from the same CATH zinc finger superfamily (CATH ID: 3.30.160.60) in the training dataset enhances the predictive accuracy, with a Pearson correlation coefficient of 0.63 and Spearman's rank correlation coefficient of 0.60.

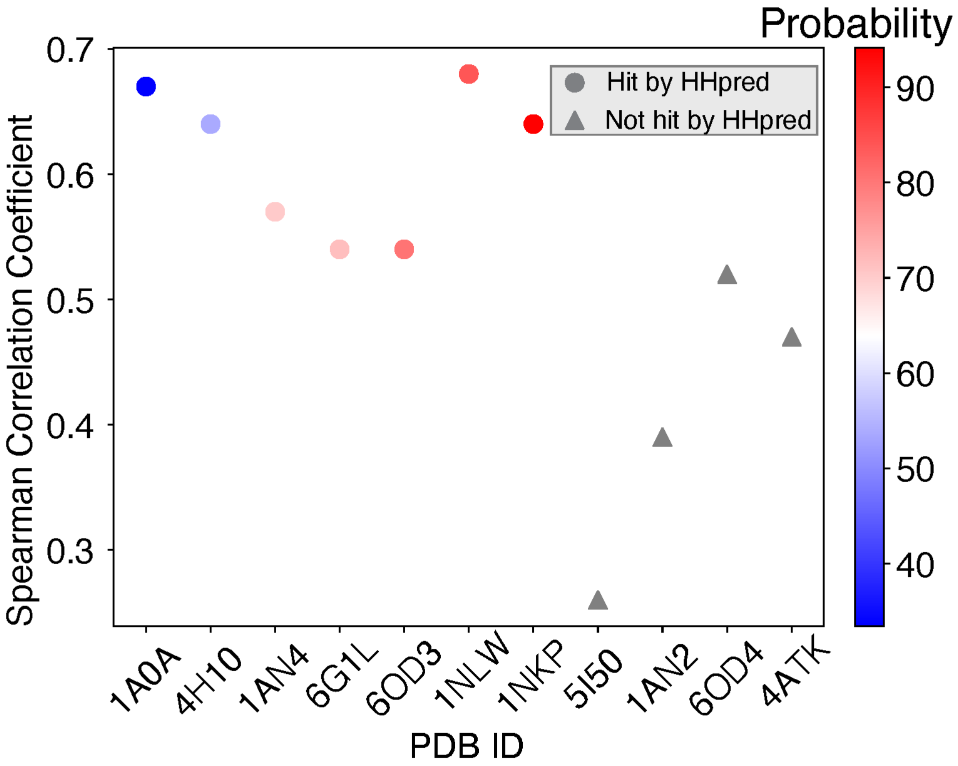


Figure S10: **Spearman's rank correlation coefficients between predicted and experimental MAX binding affinities across training proteins ordered by probability of being homologous to the MAX protein.**

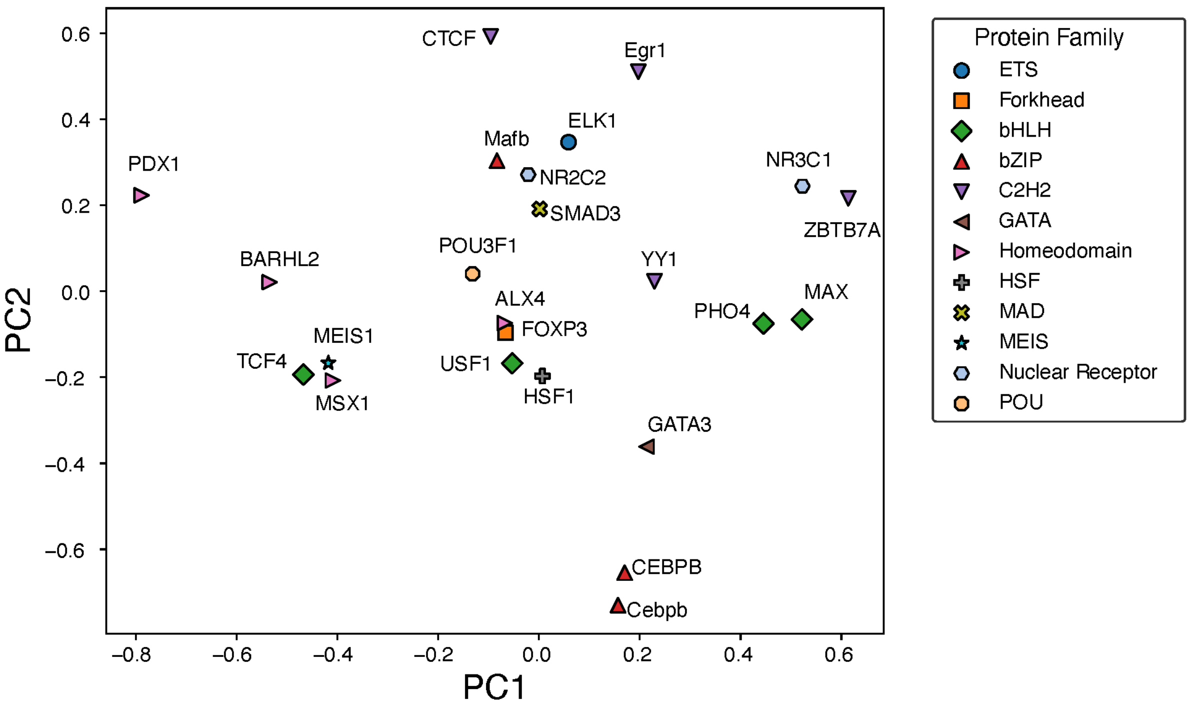


Figure S11: **Principal component analysis of the normalized IDEA-learned energy model across 12 protein families.**

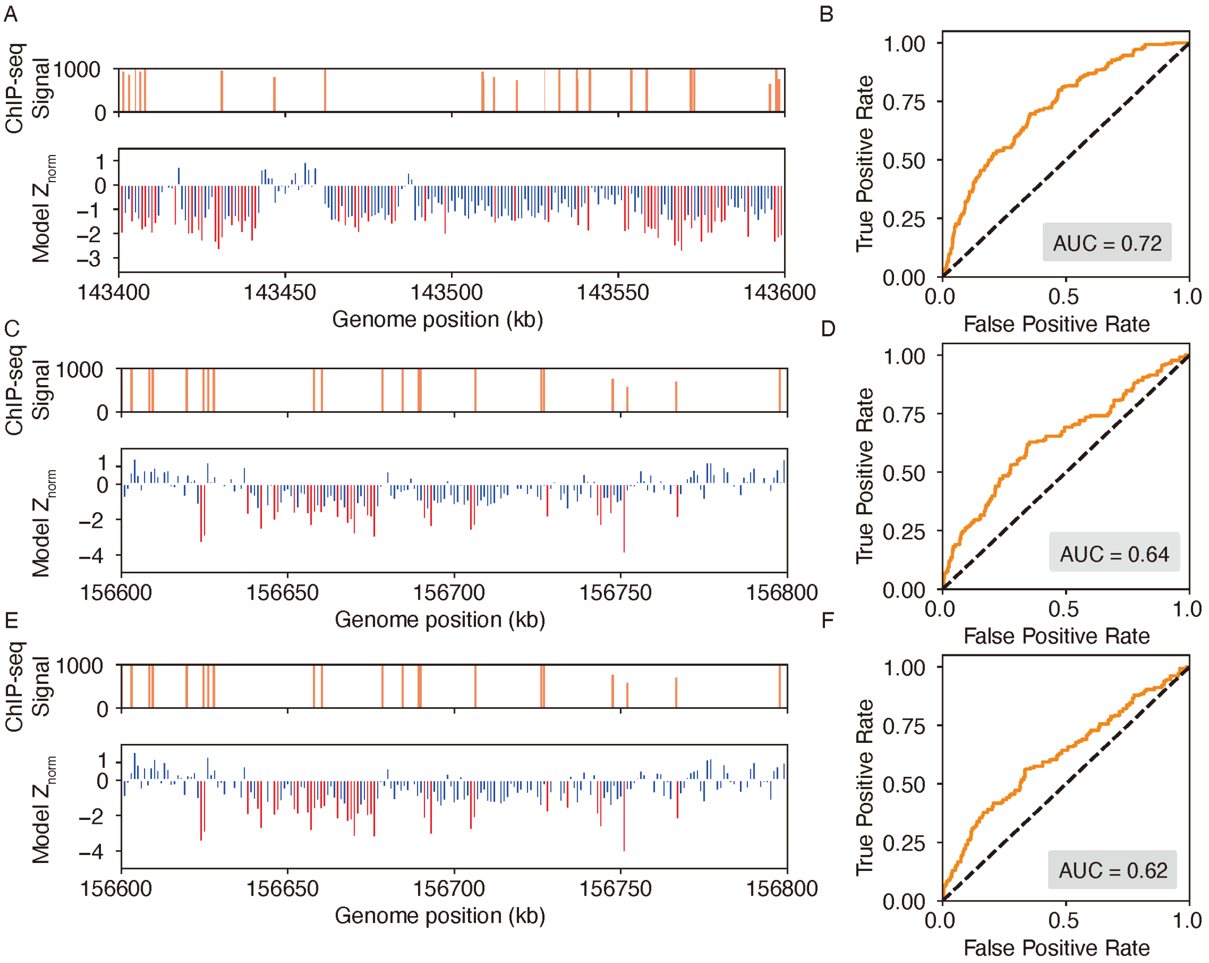


Figure S12: **IDEA accurately identifies genomic binding sites for additional proteins.** (A, B) Predicted binding sites for the EGR1 transcription factor in a 1 Mb region on chromosome 8 of the HepG2 cell line, where ChIP-seq signals are densest. The top plot of (A) shows ChIP-seq signals, while the bottom plot shows IDEA's predicted binding sites, with highly probable sites marked as red peaks. Normalized Z scores were averaged over a 500 bp window to align with the experimental resolution. The ROC curve (B) shows an AUC score of 0.72, reflecting the IDEA's accuracy for EGR1 binding sites prediction. (C–F) Predicted binding sites for the CTCF (CCCTC-Binding factor) protein in the GM12878 cell line on chromosome 1 using two different training structures: 8SSS (C, D) and 8SSQ (E, F). Panels (C) and (E) show IDEA's predictions compared to ChIP-seq signals, with highly probable binding sites highlighted as red peaks. Panels (D) and (F) present ROC curves with AUC scores of 0.64 and 0.62, respectively, indicating IDEA's predictive performance using each training structure.

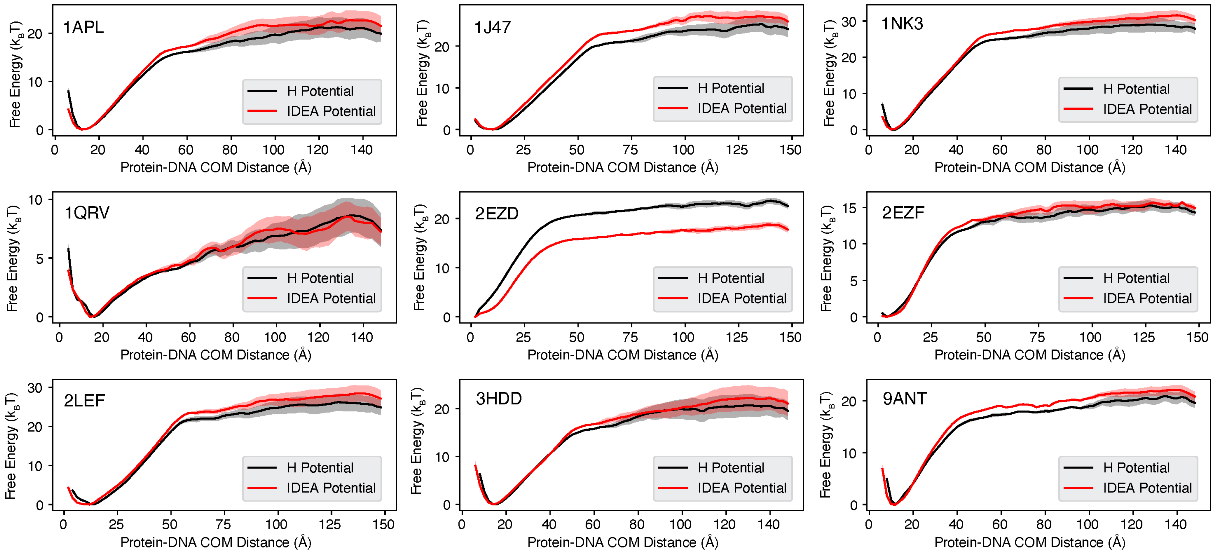


Figure S13: **Binding free energy curves calculated from simulations based on non-sequence-specific homogenous electrostatic potential and IDEA potential models.** The predicted free energy profiles were computed as a function of the Protein-DNA center of mass (COM) distance, comparing the non-sequence-specific homogenous electrostatic potential (H Potential) model (black) and the IDEA potential model (red). The line represents the mean free energy calculated from three equal partitions of the simulation trajectories, and the shaded areas were calculated as the standard deviation.

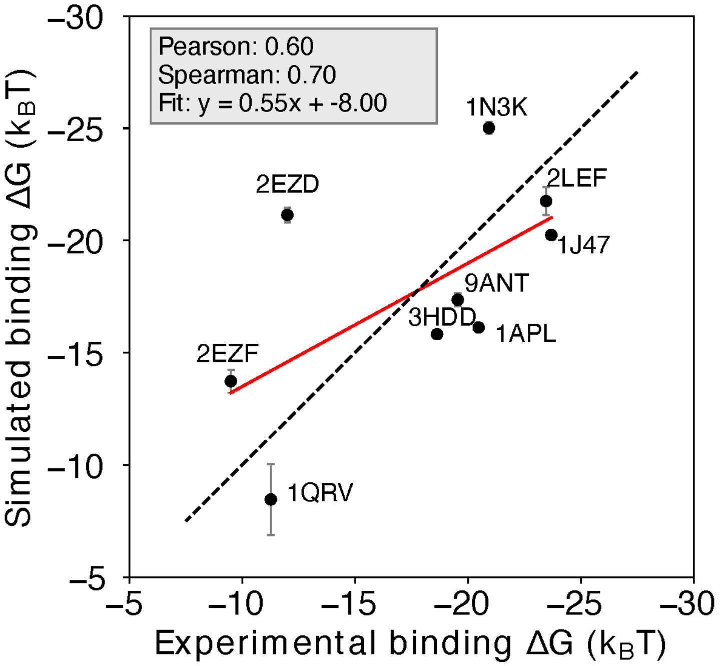


Figure S14: **Prediction of protein-DNA binding free energy with the non-sequence-specific homogeneous electrostatic potential model.** Comparison between the simulation-predicted and experimentally-assessed binding free energies for 9 protein-DNA complexes. The simulation free energies were computed using a non-sequence-specific homogenous electrostatic potential (H potential) model. The predicted binding free energies are presented in physical units. Error bars represent the standard deviation of the mean. See Figure S13 for additional details.

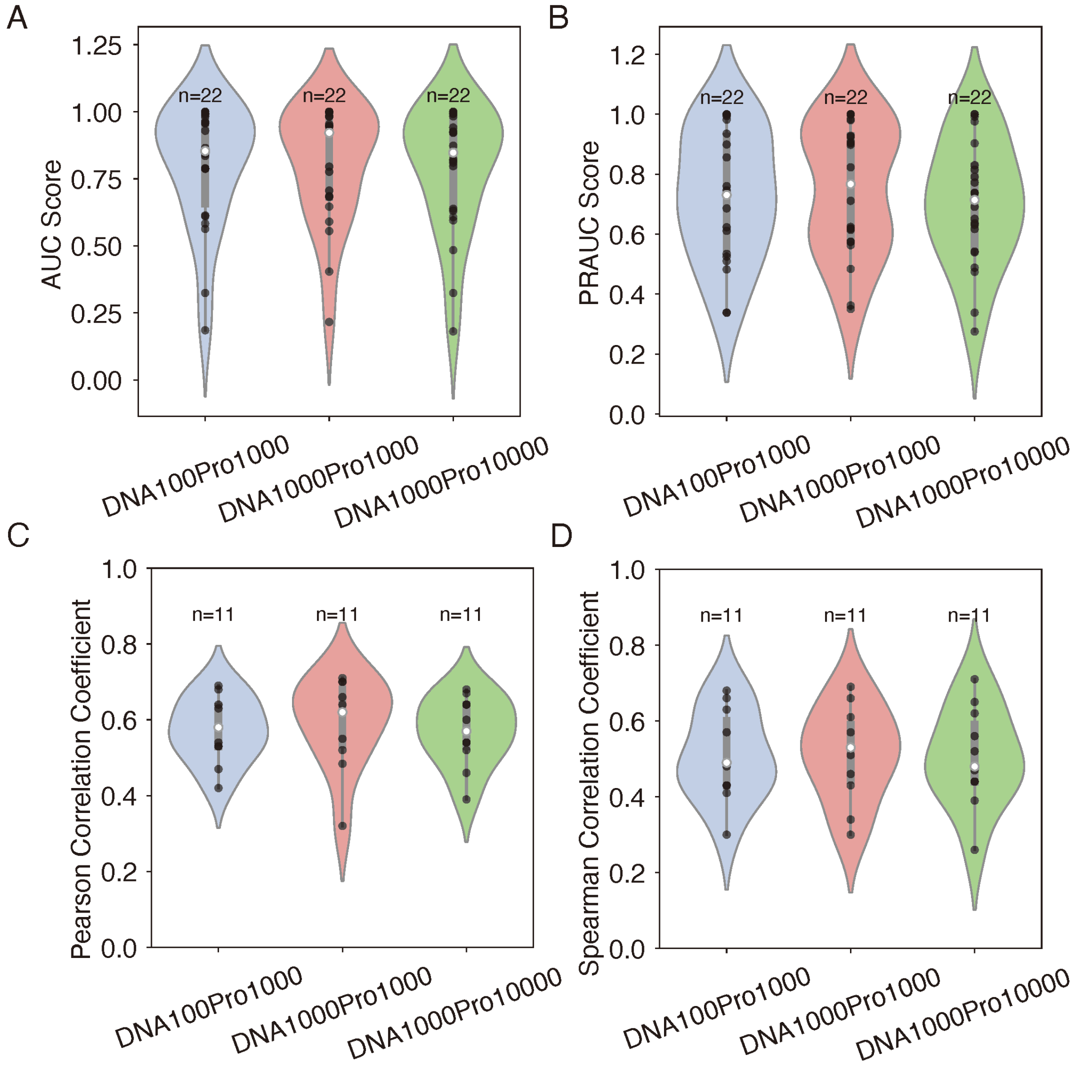


Figure S15: **Effect of the number of decoy sequences on model generalizability and transferrability.** (A-B) Evaluation of IDEA's generalizability across different decoy numbers, corresponding to Figure 2E. Violin plots summarize the distribution of AUC (A) and PRAUC (B) scores for three decoy combinations. (C-D) Analysis of IDEA transferability within the MAX CATH superfamily, corresponding to Figure 3A. Violin plots summarize the distribution of Pearson correlation coefficients (C) and Spearman's rank correlation coefficients (D) between predicted and experimental binding affinities for three decoy combinations. In each violin plot, the thick grey bar represents the interquartile range (first to third quartiles), and the thin line extends to 1.5 times the interquartile range. Individual data are depicted as scattered black points, and the white dot represents the median. Sample sizes (n) are labeled above each group. The three tested decoy combinations include: 100 DNA + 1000 protein decoys (DNA100Pro1000), 1000 DNA + 1000 protein decoys (DNA1000Pro1000), and 1000 DNA + 10000 protein decoys (DNA1000Pro10000). The consistent performance across all decoy combinations demonstrates the robustness of the model's prediction with respect to the number of decoy sequences.

### 5 Supplementary Tables

Table S1: Raw AUC scores for distinguishing strong and weak binders across 22 proteins from 12 protein families using six different prediction methods: IDEA, IDEA augmented with binding sequences (IDEA-seq), ProBound,[S10] DeepBind,[S11] DBD-Hunter[S17] and rCLAMPS[S22]

| ID | IDEA | IDEA-seq | ProBound | DeepBind | DBD-Hunter | rCLAMPS |
|---|---|---|---|---|---|---|
| ELK1 | 0.810 | 0.745 | 0.998 | 1.000 | 0.626 | – |
| FOXP3 | 0.873 | 1.000 | 0.342 | 0.812 | 0.162 | – |
| MAX | 0.941 | 1.000 | 0.736 | 0.995 | 0.490 | – |
| TCF4 | 0.638 | 1.000 | 0.827 | 0.842 | 0.090 | – |
| USF1 | 0.484 | 1.000 | 1.000 | 0.978 | 0.623 | – |
| Cebpb | 0.983 | 1.000 | 0.934 | 1.000 | 0.509 | – |
| CEBPB | 0.923 | 1.000 | 0.362 | 1.000 | 0.357 | – |
| Mafb | 0.607 | 0.987 | 0.944 | 0.998 | 0.794 | – |
| Egr1 | 0.181 | 0.857 | 0.106 | 0.983 | 0.319 | 0.214 |
| YY1 | 1.000 | 1.000 | 0.500 | 1.000 | 0.778 | 0.667 |
| ZBTB7A | 0.998 | 1.000 | 0.890 | 0.764 | 0.548 | – |
| GATA3 | 0.816 | 0.933 | 0.870 | 0.896 | 0.488 | – |
| ALX4 | 0.631 | 1.000 | 0.998 | 1.000 | 0.100 | 0.168 |
| BARHL2 | 0.822 | 0.999 | 0.934 | 0.987 | 0.723 | 0.533 |
| MSX1 | 0.926 | 0.998 | 0.732 | 0.829 | 0.382 | 0.113 |
| PDX1 | 0.595 | 0.832 | 0.719 | 0.997 | 0.539 | 0.645 |
| HSF1 | 0.993 | 1.000 | 1.000 | 1.000 | 0.742 | – |
| SMAD3 | 0.992 | 0.999 | 1.000 | 1.000 | 0.402 | – |
| MEIS1 | 0.797 | 1.000 | 0.256 | 1.000 | 0.245 | – |
| NR2C2 | 1.000 | 1.000 | 0.203 | 1.000 | 0.054 | – |
| NR3C1 | 0.921 | 0.999 | 0.539 | 1.000 | 0.724 | – |
| POU3F1 | 0.324 | 0.989 | – | 0.956 | 0.077 | – |

Table S2: Raw PRAUC scores for distinguishing strong and weak binders across 22 proteins from 12 protein families using six different prediction methods: IDEA, IDEA augmented with binding sequences (IDEA-seq), ProBound,[S10] DeepBind,[S11] DBD-Hunter[S17] and rCLAMPS[S22]

| ID | IDEA | IDEA-seq | ProBound | DeepBind | DBD-Hunter | rCLAMPS |
|---|---|---|---|---|---|---|
| ELK1 | 0.651 | 0.604 | 0.997 | 1.000 | 0.509 | – |
| FOXP3 | 0.738 | 1.000 | 0.524 | 0.795 | 0.335 | – |
| MAX | 0.772 | 1.000 | 0.556 | 0.994 | 0.449 | – |
| TCF4 | 0.617 | 1.000 | 0.796 | 0.807 | 0.317 | – |
| USF1 | 0.474 | 1.000 | 1.000 | 0.977 | 0.532 | – |
| Cebpb | 0.976 | 1.000 | 0.868 | 1.000 | 0.452 | – |
| CEBPB | 0.691 | 1.000 | 0.395 | 1.000 | 0.377 | – |
| Mafb | 0.539 | 0.985 | 0.844 | 0.998 | 0.694 | – |
| Egr1 | 0.275 | 0.746 | 0.318 | 0.980 | 0.373 | 0.345 |
| YY1 | 1.000 | 1.000 | 0.435 | 1.000 | 0.592 | 0.512 |
| ZBTB7A | 0.830 | 1.000 | 0.709 | 0.573 | 0.449 | – |
| GATA3 | 0.636 | 0.925 | 0.895 | 0.899 | 0.436 | – |
| ALX4 | 0.542 | 1.000 | 0.911 | 1.000 | 0.322 | 0.334 |
| BARHL2 | 0.737 | 0.934 | 0.434 | 0.986 | 0.598 | 0.466 |
| MSX1 | 0.815 | 0.998 | 0.584 | 0.878 | 0.401 | 0.326 |
| PDX1 | 0.488 | 0.765 | 0.540 | 0.996 | 0.455 | 0.508 |
| HSF1 | 0.992 | 1.000 | 1.000 | 1.000 | 0.580 | – |
| SMAD3 | 0.903 | 0.915 | 1.000 | 1.000 | 0.400 | – |
| MEIS1 | 0.631 | 1.000 | 0.369 | 1.000 | 0.352 | – |
| NR2C2 | 1.000 | 1.000 | 0.339 | 1.000 | 0.310 | – |
| NR3C1 | 0.791 | 0.999 | 0.513 | 1.000 | 0.586 | – |
| POU3F1 | 0.338 | 0.945 | – | 0.960 | 0.282 | – |

Table S3: This table reports the number of atoms within the cutoff distances from the C$\alpha$ atoms for all the protein-DNA structures used in this study. For each structure, we selected the DNA atom type (C5 or P) with the largest sum of occurrences within 8 Å, 9 Å, and 10 Å distances from surrounding C$\alpha$ atoms for modeling. For protein YY1 (PDB ID: 1ubd), where the counts for C5 and P were identical, both models trained on C5 and P achieved an AUC score and PRAUC score of 1.0 for distinguishing strong from weak binders of the HT-SELEX dataset (Figure S4). The $R^2$ values for 10-fold cross-validation on the HT-SELEX dataset were 0.68 for the C5-trained model and 0.647 for the P-trained model. Figures S4, S5, and S7 use the result from the C5-trained model for YY1.

| | 8 Å | 9 Å | 10 Å | Sum |
|---|---|---|---|---|
| 1a0a (C5) | 15 | 22 | 22 | 59 |
| 1a0a (P) | 18 | 19 | 20 | 57 |
| 1aay (C5) | 11 | 15 | 16 | 42 |
| 1aay (P) | 14 | 15 | 15 | 44 |
| 1an4 (C5) | 6 | 11 | 16 | 33 |
| 1an4 (P) | 16 | 18 | 24 | 58 |
| 1apl (C5) | 7 | 11 | 14 | 32 |
| 1apl (P) | 10 | 14 | 15 | 39 |
| 1dux (C5) | 8 | 9 | 11 | 28 |
| 1dux (P) | 6 | 7 | 12 | 25 |
| 1gu4 (C5) | 14 | 18 | 20 | 52 |
| 1gu4 (P) | 13 | 16 | 19 | 48 |
| 1hlo (C5) | 11 | 15 | 19 | 45 |
| 1hlo (P) | 13 | 14 | 15 | 42 |
| 1ig7 (C5) | 6 | 8 | 11 | 25 |
| 1ig7 (P) | 9 | 12 | 13 | 34 |
| 1j47 (C5) | 9 | 12 | 16 | 37 |
| 1j47 (P) | 16 | 18 | 18 | 52 |
| 1nk3 (C5) | 5 | 9 | 11 | 25 |
| 1nk3 (P) | 8 | 9 | 10 | 27 |
| 1nkp (C5) | 11 | 16 | 20 | 47 |
| 1nkp (P) | 16 | 20 | 20 | 56 |
| 1nlw (C5) | 11 | 16 | 20 | 47 |
| 1nlw (P) | 14 | 19 | 20 | 53 |

| | 8 Å | 9 Å | 10 Å | Sum |
|---|---|---|---|---|
| 1ozj (C5) | 9 | 13 | 21 | 43 |
| 1ozj (P) | 13 | 18 | 20 | 51 |
| 1qrv (C5) | 5 | 9 | 13 | 27 |
| 1qrv (P) | 9 | 9 | 11 | 29 |
| 1ubd (C5) | 14 | 21 | 24 | 59 |
| 1ubd (P) | 17 | 19 | 23 | 59 |
| 2ezd (C5) | 6 | 9 | 11 | 26 |
| 2ezd (P) | 10 | 11 | 11 | 32 |
| 2ezf (C5) | 7 | 9 | 10 | 26 |
| 2ezf (P) | 6 | 7 | 10 | 23 |
| 2h1k (C5) | 6 | 8 | 10 | 24 |
| 2h1k (P) | 8 | 11 | 12 | 31 |
| 2lef (C5) | 12 | 18 | 21 | 51 |
| 2lef (P) | 20 | 20 | 21 | 61 |
| 2wty (C5) | 13 | 19 | 23 | 55 |
| 2wty (P) | 17 | 20 | 20 | 57 |
| 2xsd (C5) | 11 | 14 | 14 | 39 |
| 2xsd (P) | 10 | 11 | 12 | 33 |
| 3hdd (C5) | 5 | 7 | 10 | 22 |
| 3hdd (P) | 7 | 8 | 11 | 26 |
| 4hc7 (C5) | 5 | 10 | 12 | 27 |
| 4hc7 (P) | 8 | 11 | 13 | 32 |
| 4xrs (C5) | 5 | 11 | 17 | 33 |
| 4xrs (P) | 11 | 14 | 15 | 40 |
| 4y60 (C5) | 7 | 11 | 14 | 32 |
| 4y60 (P) | 15 | 15 | 17 | 47 |

| | 8 Å | 9 Å | 10 Å | Sum |
|---|---|---|---|---|
| 5cbx (C5) | 8 | 15 | 17 | 40 |
| 5cbx (P) | 14 | 14 | 15 | 43 |
| 6od3 (C5) | 5 | 8 | 10 | 23 |
| 6od3 (P) | 8 | 11 | 11 | 30 |
| 7tdw (C5) | 6 | 9 | 10 | 25 |
| 7tdw (P) | 8 | 10 | 11 | 29 |
| 7xv6 (C5) | 9 | 16 | 20 | 45 |
| 7xv6 (P) | 14 | 16 | 18 | 48 |
| 8e3d (C5) | 13 | 20 | 23 | 56 |
| 8e3d (P) | 20 | 22 | 24 | 66 |
| 8k8d (C5) | 11 | 16 | 18 | 45 |
| 8k8d (P) | 9 | 14 | 16 | 39 |
| 8osb (C5) | 13 | 22 | 27 | 62 |
| 8osb (P) | 26 | 30 | 34 | 90 |
| 8pm7 (C5) | 4 | 8 | 10 | 22 |
| 8pm7 (P) | 9 | 13 | 13 | 35 |
| 8ssq (C5) | 28 | 41 | 46 | 115 |
| 8ssq (P) | 34 | 41 | 46 | 121 |
| 8sss (C5) | 29 | 39 | 42 | 110 |
| 8sss (P) | 28 | 35 | 35 | 98 |
| 9ant (C5) | 5 | 7 | 11 | 23 |
| 9ant (P) | 8 | 10 | 12 | 30 |

Table S4: Summary of parameters used in our protein-DNA simulation model (unit: kcal/mol).

| | PP | SU | A | G | C | T |
|---|---|---|---|---|---|---|
| ALA | 0 | 0 | -0.0228 | 0.0101 | 0.0199 | -0.0061 |
| ARG | 0 | 0 | -0.0402 | 0.0340 | -0.0363 | -0.0210 |
| ASN | 0 | 0 | -0.0163 | 0.0252 | 0.0159 | -0.0030 |
| ASP | 0 | 0 | -0.0001 | 0.0120 | 0.0048 | 0.0101 |
| CYS | 0 | 0 | 0.0025 | 0.0107 | 0.0062 | 0.0075 |
| GLU | 0 | 0 | 0.0045 | -0.0210 | 0.0189 | -0.0059 |
| GLN | 0 | 0 | -0.0138 | -0.0060 | 0.0094 | -0.0032 |
| GLY | 0 | 0 | 0.0027 | 0.0320 | 0.0183 | -0.0486 |
| HIS | 0 | 0 | -0.0030 | 0.0086 | 0.0149 | -0.0022 |
| ILE | 0 | 0 | -0.0188 | 0.0052 | 0.0140 | 0.0099 |
| LEU | 0 | 0 | -0.0148 | -0.0216 | 0.0127 | 0.0121 |
| LYS | 0 | 0 | -0.0107 | 0.0083 | -0.0487 | 0.0115 |
| MET | 0 | 0 | -0.0049 | 0.0085 | 0.0020 | 0.0143 |
| PHE | 0 | 0 | -0.0015 | -0.0117 | 0.0151 | -0.0042 |
| PRO | 0 | 0 | 0.0009 | 0.0240 | -0.0073 | -0.0087 |
| SER | 0 | 0 | 0.0120 | -0.0100 | 0.0011 | -0.0079 |
| THR | 0 | 0 | -0.0213 | -0.0132 | 0.0163 | 0.0066 |
| TRP | 0 | 0 | -0.0181 | 0.0058 | 0.0172 | 0.0030 |
| TYR | 0 | 0 | -0.0094 | 0.0016 | -0.0117 | 0.0127 |
| VAL | 0 | 0 | -0.0060 | 0.0102 | 0.0013 | 0.0060 |

(S1) Thompson, A. P.; Aktulga, H. M.; Berger, R.; Bolintineanu, D. S.; Brown, W. M.; Crozier, P. S.; In 'T Veld, P. J.; Kohlmeyer, A.; Moore, S. G.; Nguyen, T. D.; Shan, R.; Stevens, M. J.; Tranchida, J.; Trott, C.; Plimpton, S. J. LAMMPS - a flexible simulation tool for particle-based materials modeling at the atomic, meso, and continuum scales. *Computer Physics Communications* **2022**, *271*, 108171.

(S2) Torrie, G.; Valleau, J. Nonphysical sampling distributions in Monte Carlo free-energy estimation: Umbrella sampling. *Journal of Computational Physics* **1977**, *23*, 187–199.

(S3) Dragan, A. I.; Klass, J.; Read, C.; Churchill, M. E.; Crane-Robinson, C.; Privalov, P. L. DNA Binding of a Non-sequence-specific HMG-D Protein is Entropy Driven with a Substantial Non-electrostatic Contribution. *Journal of Molecular Biology* **2003**, *331*, 795–813.

(S4) Dragan, A. I.; Read, C. M.; Makeyeva, E. N.; Milgotina, E. I.; Churchill, M. E.; Crane-Robinson, C.; Privalov, P. L. DNA Binding and Bending by HMG Boxes: Energetic Determinants of Specificity. *Journal of Molecular Biology* **2004**, *343*, 371–393.

(S5) Dragan, A. I.; Liggins, J. R.; Crane-Robinson, C.; Privalov, P. L. The Energetics of Specific Binding of AT-hooks from HMGA1 to Target DNA. *Journal of Molecular Biology* **2003**, *327*, 393–411.

(S6) Dragan, A. I.; Li, Z.; Makeyeva, E. N.; Milgotina, E. I.; Liu, Y.; Crane-Robinson, C.; Privalov, P. L. Forces Driving the Binding of Homeodomains to DNA. *Biochemistry* **2006**, *45*, 141–151.

(S7) Privalov, P. L.; Dragan, A. I.; Crane-Robinson, C. Interpreting protein/DNA interactions: distinguishing specific from non-specific and electrostatic from non-electrostatic components. *Nucleic Acids Research* **2011**, *39*, 2483–2491.

(S8) Kumar, S.; Rosenberg, J. M.; Bouzida, D.; Swendsen, R. H.; Kollman, P. A. THE weighted histogram analysis method for free-energy calculations on biomolecules. I. The method. *J Comput Chem* **1992**, *13*, 1011–1021.

(S9) Yang, L.; Orenstein, Y.; Jolma, A.; Yin, Y.; Taipale, J.; Shamir, R.; Rohs, R. Transcription factor family-specific DNA shape readout revealed by quantitative specificity models. *Mol Syst Biol* **2017**, *13*, 910.

(S10) Rube, H. T.; Rastogi, C.; Feng, S.; Kribelbauer, J. F.; Li, A.; Becerra, B.; Melo, L. A. N.; Do, B. V.; Li, X.; Adam, H. H.; Shah, N. H.; Mann, R. S.; Bussemaker, H. J. Prediction of protein–ligand binding affinity from sequencing data with interpretable machine learning. *Nat Biotechnol* **2022**, *40*, 1520–1527.

(S11) Alipanahi, B.; Delong, A.; Weirauch, M. T.; Frey, B. J. Predicting the sequence specificities of DNA- and RNA-binding proteins by deep learning. *Nat Biotechnol* **2015**, *33*, 831–838.

(S12) Khan, A.; Fornes, O.; Stigliani, A.; Gheorghe, M.; Castro-Mondragon, J. A.; van der Lee, R.; Bessy, A.; Chèneby, J.; Kulkarni, S. R.; Tan, G.; Baranasic, D.; Arenillas, D. J.; Sandelin, A.; Vandepoele, K.; Lenhard, B.; Ballester, B.; Wasserman, W. W.; Parcy, F.; Mathelier, A. JASPAR 2018: update of the open-access database of transcription factor binding profiles and its web framework. *Nucleic Acids Research* **2018**, *46*, D260–D266.

(S13) Kulakovskiy, I. V.; Vorontsov, I. E.; Yevshin, I. S.; Sharipov, R. N.; Fedorova, A. D.; Rumynskiy, E. I.; Medvedeva, Y. A.; Magana-Mora, A.; Bajic, V. B.; Papatsenko, D. A.; Kolpakov, F. A.; Makeev, V. J. HOCOMOCO: towards a complete collection of transcription factor binding models for human and mouse via large-scale ChIP-Seq analysis. *Nucleic Acids Res* **2018**, *46*, D252–D259.

(S14) Jolma, A.; Yan, J.; Whitington, T.; Toivonen, J.; Nitta, K.; Rastas, P.; Morgunova, E.; Enge, M.; Taipale, M.; Wei, G.; Palin, K.; Vaquerizas, J.; Vincentelli, R.; Luscombe, N.; Hughes, T.; Lemaire, P.; Ukkonen, E.; Kivioja, T.; Taipale, J. DNA-Binding Specificities of Human Transcription Factors. *Cell* **2013**, *152*, 327–339.

(S15) Asif, M.; Orenstein, Y. DeepSELEX: inferring DNA-binding preferences from HT-SELEX data using multi-class CNNs. *Bioinformatics* **2020**, *36*, i634–i642.

(S16) Saito, T.; Rehmsmeier, M. The Precision-Recall Plot Is More Informative than the ROC Plot When Evaluating Binary Classifiers on Imbalanced Datasets. *PLoS ONE* **2015**, *10*, e0118432.

(S17) Gao, M.; Skolnick, J. DBD-Hunter: a knowledge-based method for the prediction of DNA-protein interactions. *Nucleic Acids Res* **2008**, *36*, 3978–3992.

(S18) Geertz, M.; Shore, D.; Maerkl, S. J. Massively parallel measurements of molecular interaction kinetics on a microfluidic platform. *Proceedings of the National Academy of Sciences* **2012**, *109*, 16540–16545.

(S19) Stormo, G. D.; Zhao, Y. Determining the specificity of protein–DNA interactions. *Nat Rev Genet* **2010**, *11*, 751–760.

(S20) Rastogi, C.; Rube, H. T.; Kribelbauer, J. F.; Crocker, J.; Loker, R. E.; Martini, G. D.; Laptenko, O.; Freed-Pastor, W. A.; Prives, C.; Stern, D. L.; Mann, R. S.; Bussemaker, H. J. Accurate and sensitive quantification of protein-DNA binding affinity. *Proc. Natl. Acad. Sci. U.S.A.* **2018**, *115*.

(S21) Maerkl, S. J.; Quake, S. R. A Systems Approach to Measuring the Binding Energy Landscapes of Transcription Factors. *Science* **2007**, *315*, 233–237.

(S22) Wetzel, J. L.; Zhang, K.; Singh, M. Learning probabilistic protein-DNA recognition codes from DNA-binding specificities using structural mappings. *Genome Res* **2022**, *32*, 1776–1786.
